# Pseudocritical and Precritical States in Brain Dynamics

**DOI:** 10.1101/2021.07.04.451067

**Authors:** Lei Gu, Ruqian Wu

## Abstract

Scale-free brain dynamics under external stimuli raises an apparent paradox since the critical point of the brain dynamics locates at the limit of zero external drive. Here, we demonstrate that relaxation of the membrane potential removes the critical point but facilitates scale-free dynamics in the presence of strong external stimuli. These findings feature biological neural networks as systems that have no real critical point but bear critical-like behaviors. Attainment of such pseudocritical states relies on processing neurons into a precritical state where they are made readily activatable. We discuss supportive signatures in existing experimental observations and advise new ones for these intriguing properties. These newly revealed repertoires of neural states call for reexamination of brain’s working states and open fresh avenues for the investigation of critical behaviors in complex dynamical systems.

## Introduction

Being the most important functional organism of living creatures, a brain needs to regularly receive and process a large amount of input signals and the correct description of its critical behavior is still under active debate [1–5]. An apparent paradox is the conflict between brain’s actual working condition and the conventional definition of the critical point of brain dynamics conferred at the limit of zero external drive [2, 6]. The criticality hypothesis is supported by increasing observations of scale-free dynamics [7–13], where the size (S) and duration (T) distributions of activity avalanches are found to follow power law statistics, *P* (*S*) ∝ *S*^−1.5^ and *P* (*T*) ∝ *T* ^−2^. The paradoxical situation and the fact that the observed exponents are in a range around the critical values (1.5, 2.0) suggest the pertinence of the quasicriticality theory [2, 14], which states that it still possible to have a susceptibility peak (albeit not divergent) in a regime close to the critical point and define a Widom line [15] (also cf. Fig. 1(a)). Around this line, the dynamics is scale-free with slightly varied exponents for brain activities.

**FIG. 1.**
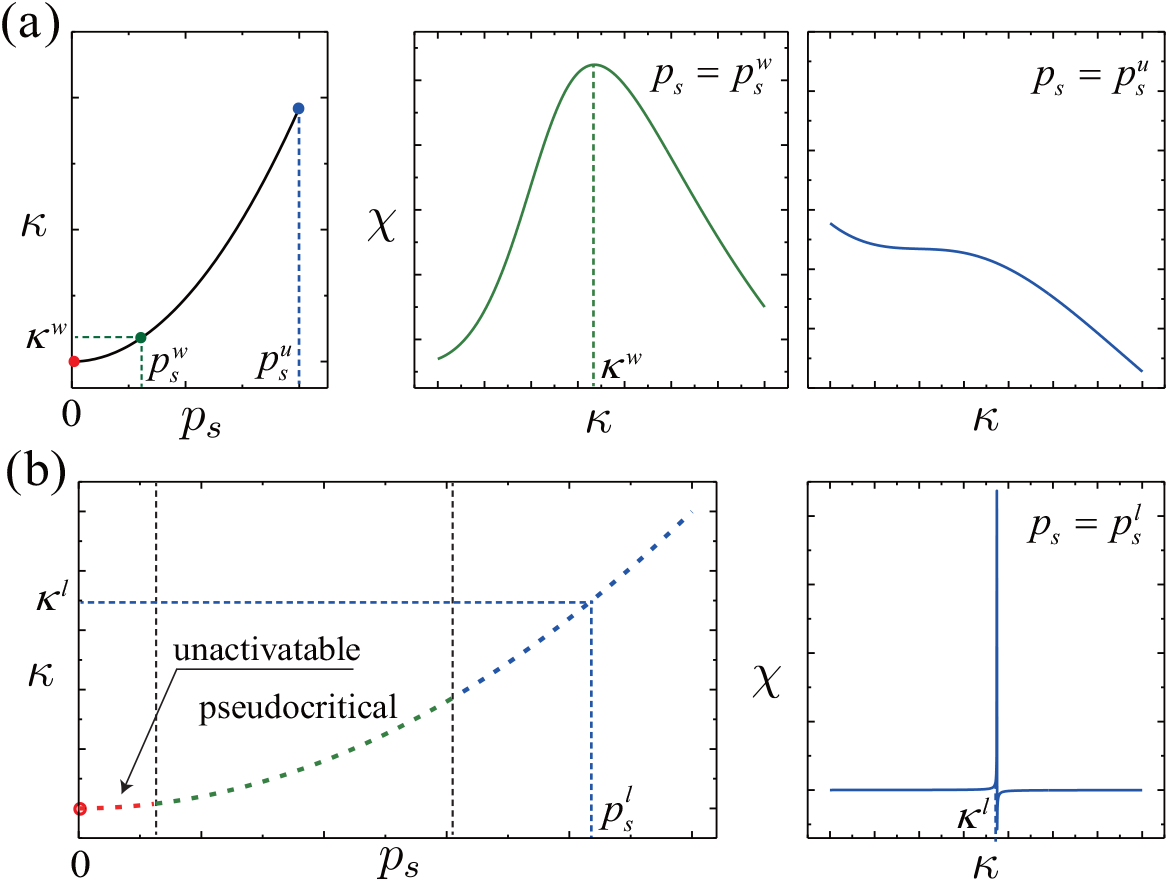
Illustration of (a) quasicriticality and (b) pseudocriticality. (a) Close to the critical point (red dot), a Widome line can be defined according to the position of susceptibility peaks until an upper bound where a susceptibility peak is not well defined; here *p*_*s*_, *κ, χ* denote frequency of external drive, branching ratio and susceptibility which will be introduced soon. (b) Under the strong external drive approximation of our proposed model, divergent susceptibility is available along a line extending from the critical point. Following a region where external stimuli (dotted red) are too weak to effectively cause neural activities, a regime around the line bears scale-free dynamics (dotted green). Because the critical point at *p*_*s*_ = 0 is inferred from strong external drive approximation, it is a pseudo one.

The quasicriticality is more ubiquitous in the study of liquid-gas phase transition [16, 17], and also presents in magnetic and strongly-correlated systems [18, 19]. In these cases, the Widom line can be roughly considered as an extrapolation of the phase boundary (e.g. the liquid-gas equilibrium line). Such a regime is occasionally referred to as “pseudocritical”. However, it might be more appropriate to reserve the terminology of “pseudocritical” for a region around a line defined by divergent susceptibility in the strong external stimulus approximation as we study the brain dynamics. Since the critical point is defined at the zero drive limit, the phase transition inferred from the strong stimulus approximation is actually unreachable. Indeed, under the weak external stimulus approximation, no critical point exits and hence the criticality is a pseudo one. Nevertheless, we find that in the pseudocritical regime the scale-free dynamics can still be well maintained. Notably, the applicable external stimuli can be significantly stronger than those for the quasicriticality, and the scale-free dynamics can take a wide range of exponents. This unveils a novel regime that is not critical but criticality-related and bears critical-like behaviors — which much enriches nonequilibrium statistics.

A typical activity circle of neurons constitutes building up of membrane potential, spiking and refractory. When incentives (external or postsynaptic) hit, the membrane potential piles up and the neuron launches a spike if a potential threshold is reached. During the refractory period, the neuron is insensitive to stimuli and eventually returns to the quiescent state of the resting membrane potential. In the interval of incentives, the membrane potential is gradually reduced by leaky of ions. Possibly because of its complexity for analytical study, this relaxation process is not considered in some typical neuron models, including the cortical branching model (CBM) that is believed to be able to describe the critical behavior of neural networks. The threshold criterion and the membrane potential leaky make it hard to induce a spike by a single incentive, and hence multiple incentives within an appropriate time span are necessary. Interestingly, unlike the sensitivity to stimuli [20] of dynamics close to (real) criticality, models imbued with leaky membrane potential usually impart states where the external drive is too weak to effectively activate the network [21] (cf. Fig. 1(b)). The activity avalanches are frustrated because the majority of neurons are deactivated by the membrane potential leaky.

To maintain the scale-free dynamics under this damping condition, neurons need to be preprocessed into states readily activatable. Our simulations suggest that critical branching processes run upon these states are still scale-free. It appears that the neural network undergoes two phases of activity — preprocessing to a precritical state and the ensuing activation of neurons described by the critical branching process. When efficient preprocessing is compromised, the size and duration of avalanches are found to be non-power law statistics, which makes an observable signature.

## Two-way cortical branching model

In the original CBM[2, 22, 23], a quiescent neuron is assigned with a probability of spiking (i.e., the incentive is exerted to transmit activity) when it receives an external or a postsynaptic incentive. After the spiking, the neuron undergoes *τ*_*r*_ refractory states and finally returns to quiescence. To improve this simplistic neuronal dynamics, we introduce prespiking states between the quiescent state and spiking state to model the membrane potential (cf. Fig.2). A spike occurs when the neuron goes through these states and pass the highest prespiking state *m*_0_. To model the potential leak, in each time step, we let a neuron in the prespiking states relax to the next lower state if no incentive arrives or the arrived incentives fail to be exerted. The relaxation constitutes another pathway to quiescence, and thus we dub it “two-way cortical branching model” (TWCBM). In addition to the parameter set of CBM, two new parameters are introduced— the number of prespiking states *τ*_*p*_ and the stride width *τ*_*s*_. The latter denotes the number of prespiking states being crossed when a transition from the quiescent state to a prespiking state or one among the prespiking states (from the low to high) are induced by incentives.

**FIG. 2.**
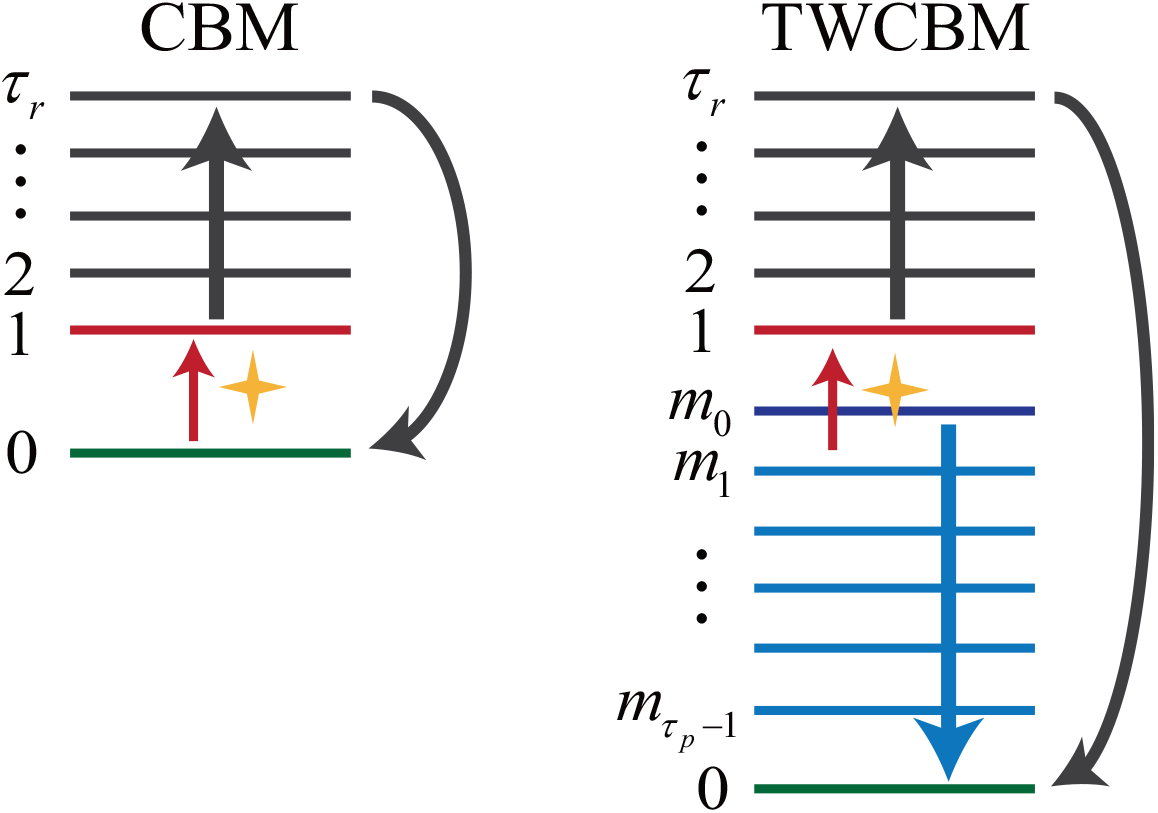
Comparison of neuronal dynamics of the CBM and TWCBM. In addition to the refractory states, prespiking states are introduced to model the membrane potential; instead of transition from the quiescent state to the first refractory state, the spiking criterion changes to the passing across the highest prespiking state (*m*_0_) and arriving at the first refractory state. The relaxation of membrane potential (blue arrow) constitutes a pathway of returning to the quiescence other than the refractory pathway (black arrow).

In the original CBM, the probability for neuron *i* to exert a postsynaptic incentive from neuron *j* takes the form *κp*_*ij*_ with ∑_*j*_ *p*_*ij*_ = 1, where *κ* is the branching ratio and the summation is over *k*_*in*_ neurons sending postsynaptic incentives to the *i*th neuron. In the TWCBM, we use this setting for strides that cross the highest prespiking state and cause spikes. For the quiescence-to-prespiking or intra-repspiking transitions, we simply set *κ* = 1 and *p*_*ij*_ = 1, that is, every incentive responsible for these transitions is exerted. As a result, the preprocessing to the precrtical states is much more efficient than the ensuing activation of neurons. After the understanding about the TWCBM is unfolded, we will discuss how this efficiency difference is implemented in realistic neurons.

## Driven to criticality

Figure 3(a) shows that, as the external stimuli are strengthened (clockwise), the network experiences states of low activity, subcritical states, (nearly) critical states, and supercritical states. Notably, weak stimuli can not effectively activate the network, and the criticality is driven by relatively strong stimuli. Ref. [24] showed that stimuli with appropriate strength can account for the decrease of the standard exponent 1.5 with strengthened stimuli. Here, we demonstrate that the scale-free dynamics can also extend to the subcritical regime and has an exponent much larger than 1.5. Since the model used there did not take into account the leak of membrane potential, it could be the key factor for the low activity and scale-free subcritical dynamics.

**FIG. 3.**
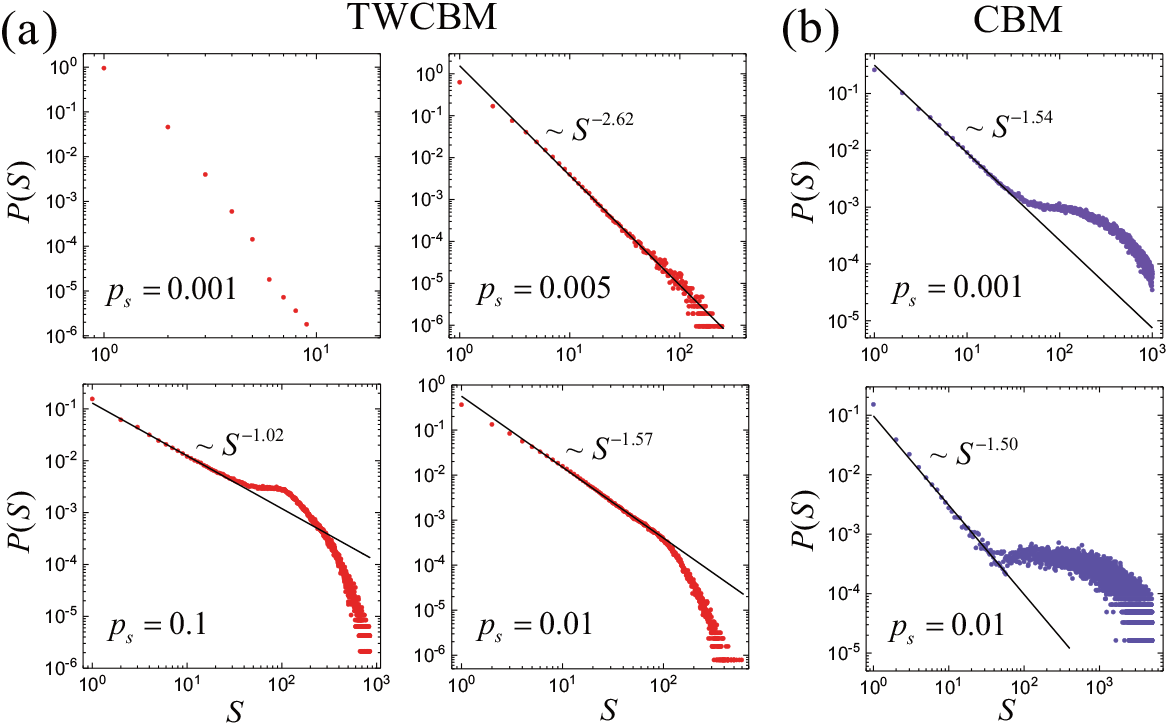
(a) Starting from the upper left and going clockwise, the TWCBM undergoes states of low activity, subcriticality, (pseudo)criticality and supercriticality. These distributions suggest that weak external stimuli cannot effectively activate the network and the dynamics is driven to the criticality only by stimuli sufficiently strong. (b) Weak stimuli can effectively activate a CBM close to criticality (upper) and significant finite size effect (the arch denoting large avalanches) suggests that the network can be easily blown up by relatively strong stimuli (lower).

To show that the TWCBM can cope with strong external stimuli better, we perform external stimuli of frequencies *p*_*s*_ = 10^*−*3^ and *p*_*s*_ = 10^*−*2^ to a CBM with the same setting (with the prespiking states excluded). As shown in the upper panel of Fig. 3(b), *p*_*s*_ = 10^*−*3^, which is too weak to activate the TWCBM, leads to dynamics close to the criticality. When the frequency increases to *p*_*s*_ = 10^*−*2^, which drives the TWCBM to the criticality, the significant finite size effect suggest that the network can be easily blown up.

In order to have correspondence with the analytical analysis carried out later, the parameters in the simulations are set as: *κ* = 1.2, *τ*_*r*_ = 5, *k*_*in*_ = 5, *τ*_*s*_ = *τ*_*p*_ = 50, and *p*_*ij*_ take biased values [2]. The network size is *N* = 128 neurons. The external stimuli are homogeneous, that is, spontaneous activation of every neuron occurs with frequency *p*_*s*_. The accompany distributions of the avalanche duration for the size distributions in this work are given in the supplementary [20].

## Pseudocritical and precritical states

Under the condition *τ*_*p*_ = *τ*_*s*_, an exerted incentive excites a neuron in the quiescent state 0 to the highest prespiking state *m*_0_, or drive a neuron in any of the prespiking states to the first refractory state and cause a spike. Therefore, in the steady state, we have rate equations

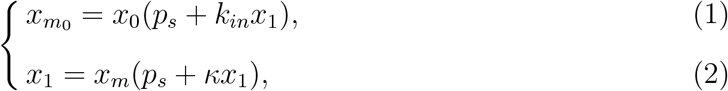

Where *x*_*m*_ denotes the total probability for a neuron to stay in the prespiking states, 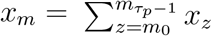. In both equations, *x*_1_ on the left hand side is the probability of a neuron being at the first refractory state, and *x*_1_ on the right hand side is the spiking rate (also the probability of generating a postsynaptic incentive) of a neuron. Because the spikes are caused by transition from the prespiking states to the first refractory state and thereafter the refractory process is unidirectional, these two probabilities are equivalent by definition. Further, because the refractory is deterministic and ergodic, *x*_*z*_ = *x*_1_ for *z* ∈ [2, *τ*_*r*_]. Here, we neglect the case of multiple incentives at a time, since *x*_1_, *p*_*s*_ ≪ 1 and the high order products are negligible. We also note that *τ*_*p*_ = *τ*_*s*_ is assumed since this enables analytical solution, but it dose not means specialized dynamics. When *τ*_*p*_ and *τ*_*s*_ are large (50 here), the setting corresponds to efficient potentiation and slow relaxation. This should be an appropriate description of the realistic situation. Otherwise, the neural dynamics may lead to barely activatable networks.

To determine spiking rate *x*_1_, two more equations are needed. One is the normalization condition

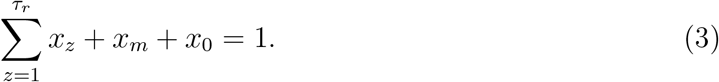

The probability of being at a prespiking state *m*_*n*_ amounts to the probability of no incentive being exerted after the neuron is exited to *m*_0_, i.e., nothing but the relaxation happens to the neuron within *n* time steps. Assuming that the external stimuli and spikes of neurons are independent Poisson processes, the summation rule for multiple Poisson processes gives

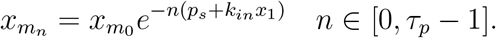

Sum of this geometric sequence gives

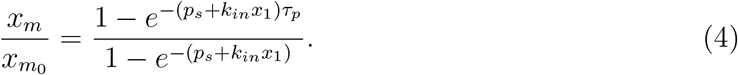

Eqs.(1-4) is sufficient to determine *x*_1_. Since Eq. (4) is a transcendental one, we solve the equations by assuming (*p*_*s*_ + *k*_*in*_*x*_1_)*τ*_*p*_ ≪ 1 and (*p*_*s*_ + *k*_*in*_*x*_1_)*τ*_*p*_ ≫ 1, that is, the weak external stimulus (and low activity) approximation and strong external stimulus (and high activity) approximation, respectively.

With the weak stimulus approximation, the solution for *p*_*s*_ = 0 is given by

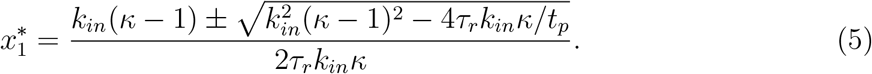

The susceptibility defined as 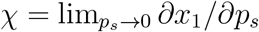 reads

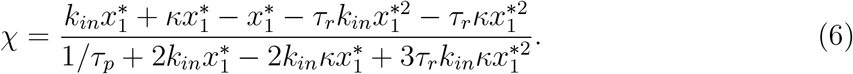

The critical point is located by finding *κ* that leads to 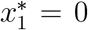 and χ → ∞ [2]. However, 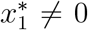 for meaningful parameter values (*τ*_*r*_, *k*_*in*_, *κ >* 0) and the denominator of χ is finite even if 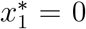. Therefore, no critical point exists under the weak stimulus approximation. This is consistent with the barely activatable network under weak external stimuli.

The strong stimulus approximation gives

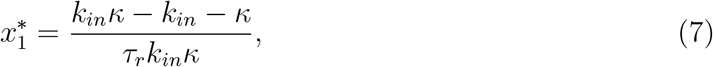

and

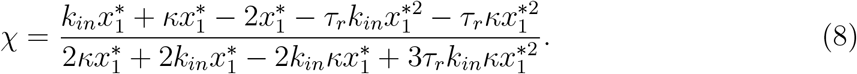

Then *κ*_*c*_ = *k*_*in*_*/*(*k*_*in*_ *−* 1) makes 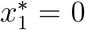 and the denominator of χ zero. Expansions around this critical value leads to 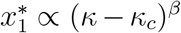 and χ ∝ (*κ − κ*_*c*_)^*−γ*^ with *β* = *γ* = 1, the same with the original CBM. While it shows critical behavior belonging to the universality class of the absorbing-state phase transitions, we note that the critical point is unreachable, since the zero drive limit contradicts with the assumption (*p*_*s*_ + *k*_*in*_*x*_1_)*t*_*p*_ ≫ 1.

When *p*_*s*_ *≠* 0, the solution is hard to derive. We numerically find the solution of Eqs.(1-4) with the same parameter setting as the simulation, and in Fig. 4 compare the curves *x*_1_(*p*_*s*_), χ(*p*_*s*_) and χ(*κ*) (with fixed *p*_*s*_) to those of the original model (insets). All these dependences are qualitatively different for TWCBM and CBM. The slow increase of *x*_1_ at small *p*_*s*_ is consistent with the absence of a critical point in the TWCBM. Close to *p*_*s*_ *≈* 0.005, *x*_1_ starts to increase rapidly. Accordingly, unlike χ of CBM that peaks at *p*_*s*_ = 0, χ of the TWCBM peaks at *p*_*s*_ *≈* 0.005. To compare the χ(*κ*) dependence in the critical regime (i.e., *P* (*S*) *≈ S*^*−*1.5^), Fig. 4(c) shows results for *p*_*s*_ = 0.01, 0.001 from the TWCBM and CBM, respectively. One can see that there is no peak for the TWCBM. This is understandable, since the precritcal states set size limit to the ensuing avalanches, and thereby high branching ratios do not lead to hyperactive networks that have low sensitivity to the variation of external drive. This monotonic increase is confirmed by numerical simulation [20].

**FIG. 4.**
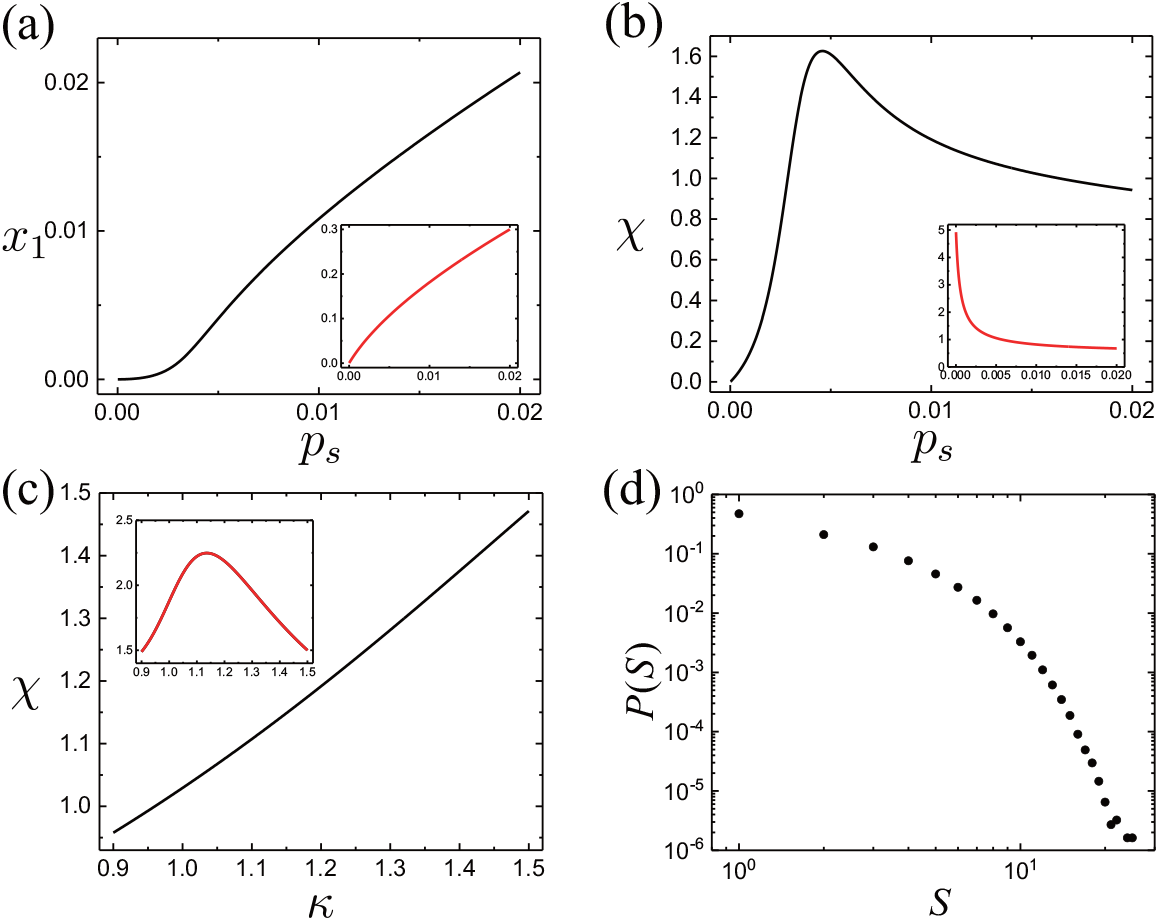
(a) Only above a certain *p*_*s*_ the activity starts to increase rapidly, which is consistent with the susceptibility peak at finite *p*_*s*_ in (b). (c) The χ(*κ*) dependence does not necessarily have a peak for dynamics in the pseudocritical regime. For comparison, the insets present corresponding properties of a nearly critical CBM. (d) If the preprocessing to precritical states is not more efficient than ensuing activation of neurons, non-power law distribution of avalanche size results as large scale precritical states are unlikely to be formed.

So far, we set the exerting rate of a postsynaptic incentive to 1 when it is responsible for excitations into the prespiking states, so that we have excitation rate *k*_*in*_*x*_1_. Since *k*_*in*_*x*_1_ *≫ κx*_1_ for *k*_*in*_ *≫* 1, this setting implies that the preprocessing to the precritical states is much more efficient than the ensuing activation of neurons. When the exerting rate is reduced, we can have the distributions in Fig. 4(d), where *k*_*in*_*x*_*l*_ is rescaled by a factor *κ*_*p*_ so that *κ*_*p*_*k*_*in*_*x*_*l*_ = *κx*_1_. Namely, we assign equal exerting rate to incentives responsible for excitations into the prespiking states and those causing spikes. The resultant distribution of avalanche size follows non-power laws (Fig. 4(d)).

In realistic neurons, the efficiency reduction in the activation phase could result from limited recovery of neurotransmitters, since it closely ensues the preprocessing which has consumed considerable amount of accumulated neurotransmitters. This observation also implies that efficient preprocessing requires activity intervals, within which the neurotransmitters can sufficiently pile up. The blurring of the efficiency difference may be the reason why homogeneous networks of the practical model proposed in Refs. [25–27] do not bear scale-free dynamics by the time-bin measure [4]. Not to digress much, we expatiate this point in the supplementary [20].

## Outlook and conclusion

In Ref. [28], it was observed that membrane potential plateaus are built up in the quiescence intervals and decline in the bursting periods. Namely, the neural tissue is processed to a highly potentiated state during the intervals and the activity bursts release the potential. This is consistent with the picture of precritical state. Ref. [12] showed that, in the regime of consecutive activity, the size distribution of avalanches appears to be non-power laws, which bear a resemblance to Fig.4(d). The effect of the blurred efficiency difference discussed above may have been observed, but taken as an ordinary subcritical signature. These experiments were carried out *in vivo*. More controllable *in vitro* measurements and careful comparison with model simulations would give more reliable support to the dynamics described by the TWBCM.

The presence of the precritical states is supposed to result in more selective response to external drive. For instance, a sequence of stimuli onto a single neuron should be less effective than distributed stimuli to multiple neurons in terms of processing the network to the precritical states, as repeated stimuli onto a single neuron may quickly activate the nearby neurons and large scale precritical states are less likely. It is desirable for a functional apparatus to be responsive to informative inputs instead of arbitrary stimuli. This selection ability may impart functional advantages aside from those entailed by the criticality [23, 29– 36]. The key dynamics ingredients rendering the precritcal and pesudocritical states are fast warming up and slow relaxation, which are not necessarily restricted to neural networks. The TWCBM can be a prototype for studies of other complex dynamical systems, especially for functional ones that need to deal with input signals.

The importance of the present study is threefold. First, we propose the two-way cortical branching model which does not change the universality class and gives a more complete description of the neural dynamics than the original model. Second, we reveal a strategy of achieving scale-free dynamics which copes with strong external drive better than vanilla branching processes. The dynamics can be viewed as critical branching processes run upon precritical states in which neurons are preprocessed to be activatable. Third, we identify a type of neural states dubbed pesudocritical states. These states cannot be extrapolated to the critical point defined at the zero drive limit, whereas they bear scale-free dynamics. In conclusion, this work puts forward biological neural networks, including the human brain, as a novel class of systems where no critical point exists but apparent criticality can occur.

This work was supported by the National Science Foundation through the University of California-Irvine Material Research Science and Engineering Center (grant no. DMR2011967). Codes for the simulations will be deposited on GitHub along with publication of this work.

